# A bireporter recombinant SARS-CoV-2 Omicron BA.5 for *in vitro* and *in vivo* studies

**DOI:** 10.64898/2026.01.24.700018

**Authors:** Esteban M. Castro, Ramya S. Barre, Chengjin Ye, Brian Imbiakha, Shahrzad Ezzatpour, Anastasija Cupic, Mark R. Walter, James J. Kobie, Richard K. Plemper, Adolfo García-Sastre, Ahmed Mostafa, Hector C. Aguilar, Luis Martinez-Sobrido

## Abstract

The continuous emergence of variants of concern (VoC) represent a significant challenge to effectively control SARS-CoV-2. Although FDA-approved vaccines and antivirals have been successfully developed and implemented for the prophylactic and therapeutic intervention of SARS-CoV-2 infection, recent VoC could escape protection garnered by previous vaccine and antiviral approaches. Determining the efficacy of prophylactics and/or therapeutics against recent VoC will assist in efficiently controlling currently circulating SARS-CoV-2 strains. We used our previously described bacterial artificial chromosome (BAC)-based reverse genetics approach for Omicron BA.5 to generate a recombinant SARS-CoV-2 BA.5 encoding a fusion of ZsGreen to Nanoluciferase (rBA.5 ZsG-Nluc) from the locus of the viral nucleocapsid (N) protein separated by the porcine teschovirus-1 (PTV-1) 2A proteolytic cleavage site. The rBA5 ZsG-Nluc replicates to levels comparable to recombinant BA.5 wild-type (rBA.5 WT) and expresses high levels of ZsG and Nluc in cultured cells. This facilitates tracking viral infection and the identification of antivirals and neutralizing antibodies (NAbs) with EC_50_ and NT_50_ values, respectively, similar to those obtained with rBA.5 WT. Importantly, in K18 hACE2 mice, rBA.5 ZsG-Nluc retains the same pathogenicity and ability to replicate in the lungs of infected mice as rBA.5 WT. Using rBA.5 ZsG-Nluc, we detected Nluc activity systemically and Nluc and ZsG expression in the lungs of infected mice using an *in vivo* imaging system (IVIS). Our results demonstrate the feasibility of using rBA.5 ZsG-Nluc to track viral infections and identify prophylactics and therapeutics against recent SARS-CoV-2 VoC *in vitro*, *ex vivo*, and *in vivo*.

**IMPORTANCE:** SARS-CoV-2, the causative virus of the COVID-19 pandemic, is continually evolving to escape immunity acquired by previous natural infections or vaccinations. Moreover, recent SARS-CoV-2 variants of concern (VoC) have acquired antiviral resistant mutations to FDA-approved drugs. The emergence of these VoC highlights the importance of identifying new prophylactics and therapeutics against currently circulating SARS-CoV-2 strains. We generated a recombinant bireporter Omicron BA.5 SARS-CoV-2 (rBA.5 ZsG-Nluc) that expresses reporter proteins, which are useful for cellular and whole animal studies and has similar viral replication and pathogenicity to a wild-type (WT) recombinant Omicron BA.5 SARS-CoV-2 (rBA.5 WT). In K18 hACE2 mice, rBA.5 ZsG-Nluc infection can be tracked systemically or in the lungs of infected mice using an *in vivo* imaging system (IVIS). We establish a proof-of-concept platform of rBA.5 ZsG-Nluc in combination with an ancestral SARS-CoV-2 strain expressing mCherry to simultaneously identify antivirals and neutralizing antibodies against original and recent SARS-CoV-2 strains.

## INTRODUCTION

Severe acute respiratory syndrome coronavirus 2 (SARS-CoV-2) is a single stranded, positive-sense, enveloped RNA virus belonging to the *Coronaviridae* family (1). SARS-CoV-2 has been responsible for the coronavirus disease 2019 (COVID-19) pandemic, resulting in 779 million infections and 7.1 million human deaths worldwide (2).

Prophylactic vaccines have proven to be effective in reducing disease severity and hospitalization in infected individuals, but breakthrough infections have led to the continued presence of SARS-CoV-2 in the population (3, 4). Moreover, SARS-CoV-2 has evolved to evade acquired immunity by previous natural infections or vaccination strategies, leading to the emergence of new viral variants, including pathogenic variants of concern (VoC) (5–9). Whereas vaccination formulations have been updated to protect against more recent viral variants, SARS-CoV-2 is still evolving and still responsible for a significant number of human infections across 6 continents (2, 10).

Antiviral therapeutics assist those who are susceptible to severe disease and/or cannot mount protective immune responses after vaccination, such as immunocompromised patients (11–13). Currently, the three major antivirals for the treatment of SARS-CoV-2 infection are Paxlovid (nirmatrelvir targeting the main viral protease or Mpro combined with nirmatrelvir metabolic inhibitor ritonavir), and two nucleoside-analog inhibitors of the viral RNA dependent RNA polymerase (RdRp), Remdesivir and Molnupiravir (14, 15). However, the use of antivirals in the clinic has resulted in the emergence of drug-resistant variants able to overcome Paxlovid and Remdesivir (16–18). These drug-resistant strains with distinct mutations emerged in infected patients who received Paxlovid or Remdesivir monotherapies, although they were in relatively low frequencies and did not increase the risk of viral spread (19).

Reverse genetic systems facilitate the generation of recombinant viruses, including those expressing reporter genes to study different aspects of their biology (18, 20–24). One of the advantages of recombinant viruses expressing reporter genes, such as those encoding fluorescent proteins, is to track and quantify viral infections *in vitro* using fluorescent microscopy or fluorescent plate readers (20, 25). Likewise, fluorescent proteins allow for the detection of viral infection *ex vivo* in tissues from infected animals using *in vivo* imaging systems (IVIS) (21, 26). However, fluorescent proteins are not well-suited for *in vivo* imaging of entire animals (20, 27). For this task, luciferase proteins represent a better option, as they allow for the detection of reporter gene expression in the entire animal using IVIS (28, 29). Likewise, luciferase proteins can be detected *ex vivo* in tissues from infected animals. However, luciferase proteins are poorly suited to identify individual infected cells (27). To overcome the limitations of either reporter technology, we have previously demonstrated that it is feasible to generate dual-reporter viruses, including SARS-CoV-2, expressing both fluorescent and luciferase (nanoluciferase, Nluc) reporter genes (20, 21, 30).

In this study, we use our previously described bacterial artificial chromosome (BAC)-based reverse genetics system for Omicron BA.5 to generate a recombinant virus expressing a fusion of a fluorescent ZsGreen and Nluc (rBA.5 ZsG-Nluc) from the locus of the viral nucleocapsid (N) protein, using a porcine teschovirus-1 (PTV-1) 2A proteolytic cleavage site approach (20, 30). The rBA.5 ZsG-Nluc replicates *in vitro* to levels comparable to the recombinant Omicron BA.5 wild-type (rBA.5 WT) and is able to stably express high levels of both reporter genes in cultured cells, allowing for viral detection and quantification of fluorescence (ZsG) and luminescence (Nluc), including the uncovering of antivirals and neutralizing antibodies (NAb) with EC_50_ and NT_50_, respectively, comparable to those obtained with rBA.5 WT. Notably, rBA.5 ZsG-Nluc retains viral pathogenicity and levels of viral replication similar to rBA.5 WT in K18 hACE2 mice. Importantly, we have assessed viral infection *in vivo* (Nluc) in the entire mouse or *ex vivo* (Nluc and ZsG) in the lungs of K18 hACE2 mice infected with rBA.5 ZsG-Nluc using IVIS. In addition, we combined a previously described recombinant SARS-CoV-2 based on the Washington-1 (WA.1) original strain expressing mCherry-Nluc (rWA.1 mCherry-Nluc) with rBA.5 ZsG-Nluc in a bireporter assay to identify antivirals and NAbs against both, the ancestral and currently circulating SARS-CoV-2 strains (20). Our results demonstrate the feasibility of using a PTV-1 2A proteolytic cleavage site to generate recombinant SARS-CoV-2 expressing a tandem reporter cassette from the locus of the N protein for a recent Omicron BA.5 strain. This rBA.5 ZsG-Nluc dual reporter strain supports identification of prophylactics and therapeutics against recent SARS-CoV-2 VoC, alone or in combination with an original SARS-CoV-2 WA.1 expressing mCherry, which afforded the simultaneous identification of antivirals or NAbs against ancestral and contemporary SARS-CoV-2 strains.

## RESULTS

### Generation and *in vitro* characterization of rBA.5 ZsG-Nluc

To generate a replication-competent recombinant SARS-CoV-2 Omicron BA.5 expressing ZsG-Nluc (rBA.5 ZsG-Nluc), a ZsG-Nluc fusion construct was inserted upstream of the viral N gene. The reporters were separated from the viral N gene by a porcine teschovirus-1 (PTV-1) 2A autoproteolytic cleavage site (**Fig. 1A**). Following viral rescue in Vero AT cells, we examined ZsG fluorescent expression by fluorescent microscopy (**Fig. 1B**). Vero AT cells were mock-infected or infected (MOI 1) with rBA.5 WT or rBA.5 ZsG-Nluc. At twelve hpi, cells were fixed and immunostained with the N-specific MAb 1C7C7, followed by a TxRed-conjugated secondary Ab and DAPI staining. Fluorescence microscopic analysis revealed ZsG signal in rBA.5 ZsG-Nluc–infected cells, which colocalized with TxRed signal indicating N protein detection. As expected, we did not observe ZsG expression in cells infected with rBA.5 WT, but we detected the presence of virus by immunostaining with the anti-N MAb. No ZsG expression or N-positive cells were observed in mock-infected cells (**Fig. 1B**). These results demonstrate that ZsG is expressed from rBA.5 ZsG-Nluc-infected cells and that ZsG expression correlates with the presence of viral N antigen.

**Figure 1.**
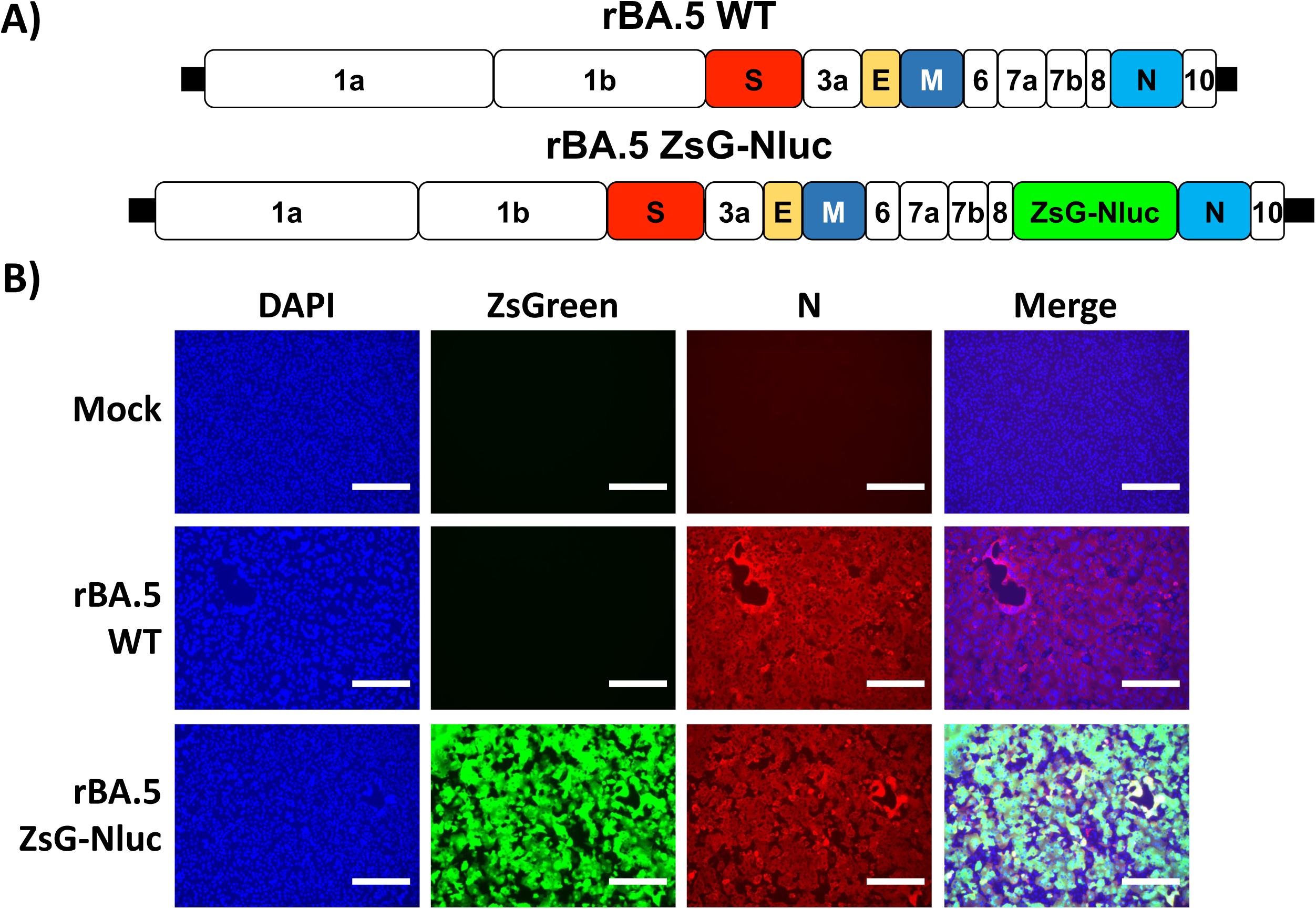
Generation and characterization of rBA.5 ZsG-Nluc. (**A**) Schematic representation of recombinant SARS-CoV-2 BA.5 wild-type (rBA.5 WT, top) and recombinant bireporter SARS-CoV-2 BA.5 (rBA.5 ZsG-Nluc, bottom). SARS-CoV-2 structural and open reading frame (ORF) accessory proteins are indicated in colors and white, respectively. (**B**) Vero AT cells were mock-infected or infected (MOI = 1) with rBA.5 WT or rBA.5 ZsG-Nluc. At 12 hpi, cells were observed under a fluorescent microscope to visualize ZsG (green), N protein (red), and nuclei (blue). Scale bar = 300 µm.

### Viral growth kinetics and plaque phenotype

To evaluate viral replication, we compared the growth kinetics of rBA.5 ZsG-Nluc to rBA.5 WT by assessing viral titers (PFU/mL) at different times post-infection and by plaque assay (20). Vero AT cells were infected (MOI 0.001) with both viruses, and CCSs were collected at 12, 48, and 72 hpi for viral titration. At these same time points, fluorescence images were taken to assess ZsG expression in both rBA.5 WT- and rBA.5 ZsG-Nluc-infected cells. Viral cytopathic effect (CPE) was evaluated by bright field imaging. CPE peaked at 48 hpi and was comparable in both rBA.5 WT and rBA.5 ZsG-Nluc infected cells (**Fig. 2A**). These results were supported by peak ZsG expression (**Fig. 2A**), Nluc activity (**Fig. 2B**), and viral titers (**Fig. 2C**) at this time point. Notably, viral titers of rBA.5 ZsG-Nluc and rBA.5 WT were comparable at all times post-infection (**Fig. 2C**). Plaque assay results show that rBA.5 ZsG-Nluc, but not rBA.5 WT, plaques expressed ZsG and Nluc (**Fig. 2D**). Crystal violet staining suggested that plaques from rBA.5 WT were slightly bigger than those of rBA.5 ZsG-Nluc (**Fig. 2D**). These results show that rSARS-CoV-2 BA.5 ZsG-Nluc expresses both reporter genes and replicates in Vero AT cells to levels comparable to rSARS-CoV-2 WT.

**Figure 2.**
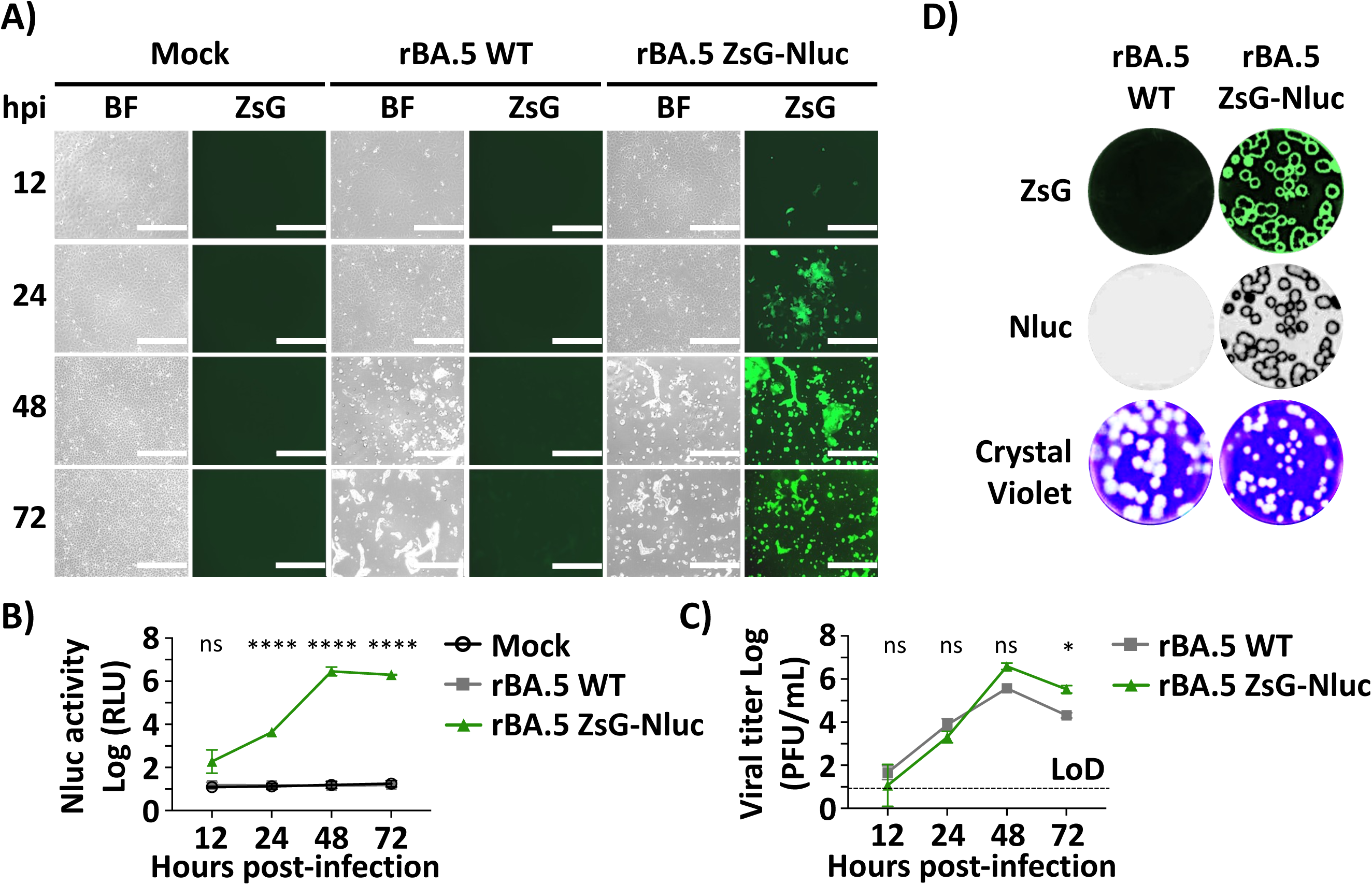
Growth kinetics and plaque phenotype of rBA.5 ZsG-Nluc. Vero AT cells were mock-infected or infected (MOI = 0.001) with rBA.5 WT or rBA.5 ZsG-Nluc and CCSs were collected at 12, 24, 48, and 72 hpi. (**A**) Brightfield (BF) and ZsG fluorescence images of cells at the indicated timepoints. Scale bar = 400 µm. (**B**) NanoLuc (Nluc) expression in CCSs collected from (**A**) measured using a PheraStar reader. (**C**) Viral titers present in the CCSs as determined by plaque assay. (**D**) Plaque morphology assessed by fluorescence (ZsG), luminescence (Nluc), and crystal violet staining at 72 hpi. Statistical analysis performed using two-way ANOVA with Geiser-Greenhouse correction and Sidak post hoc test. Dotted line in panel C indicates the limit of detection (LoD). Data are presented as mean ± SD. ****, P < 0.0001. ***, P < 0.001. ns, non-significant.

### Microneutralization assays for the identification of antivirals and NAbs

The current challenge in managing SARS-CoV-2 infections lies in the emergence of VoC able to escape vaccine-induced immune responses (5). Furthermore, the reliance on antiviral monotherapies using protease inhibitors has facilitated the emergence of drug-resistant mutations (17). Another concern is that ancestral SARS-CoV-2 strains do not accurately represent currently circulating SARS-CoV-2 VoC in terms of susceptibility to neutralizing antibodies or antivirals. We consider contemporary strains, like rBA.5 ZsG-Nluc, a superior option to assess prophylactics and therapeutics against currently circulating SARS-CoV-2 (6, 46–48).

To test feasibility of using rBA.5 ZsG-Nluc to identify antivirals, A549-hACE2 cells were infected with rBA.5 WT or rBA.5 ZsG-Nluc and then treated with 2-fold serial dilutions of the FDA-approved Mpro inhibitor drug combination Paxlovid (nirmatrelvir + ritonavir), or the RDRP inhibitor Remdesivir (**Fig. 3A**). Both rBA.5 ZsG-Nluc and rBA.5 WT were similarly inhibited by Paxlovid (EC_50_ = 0.01 µM and 0.03 µM, respectively) and Remdesivir (EC_50_ = 8.08 µM and 10.55 µM, respectively) (**Fig. 3A**).

**Figure 3.**
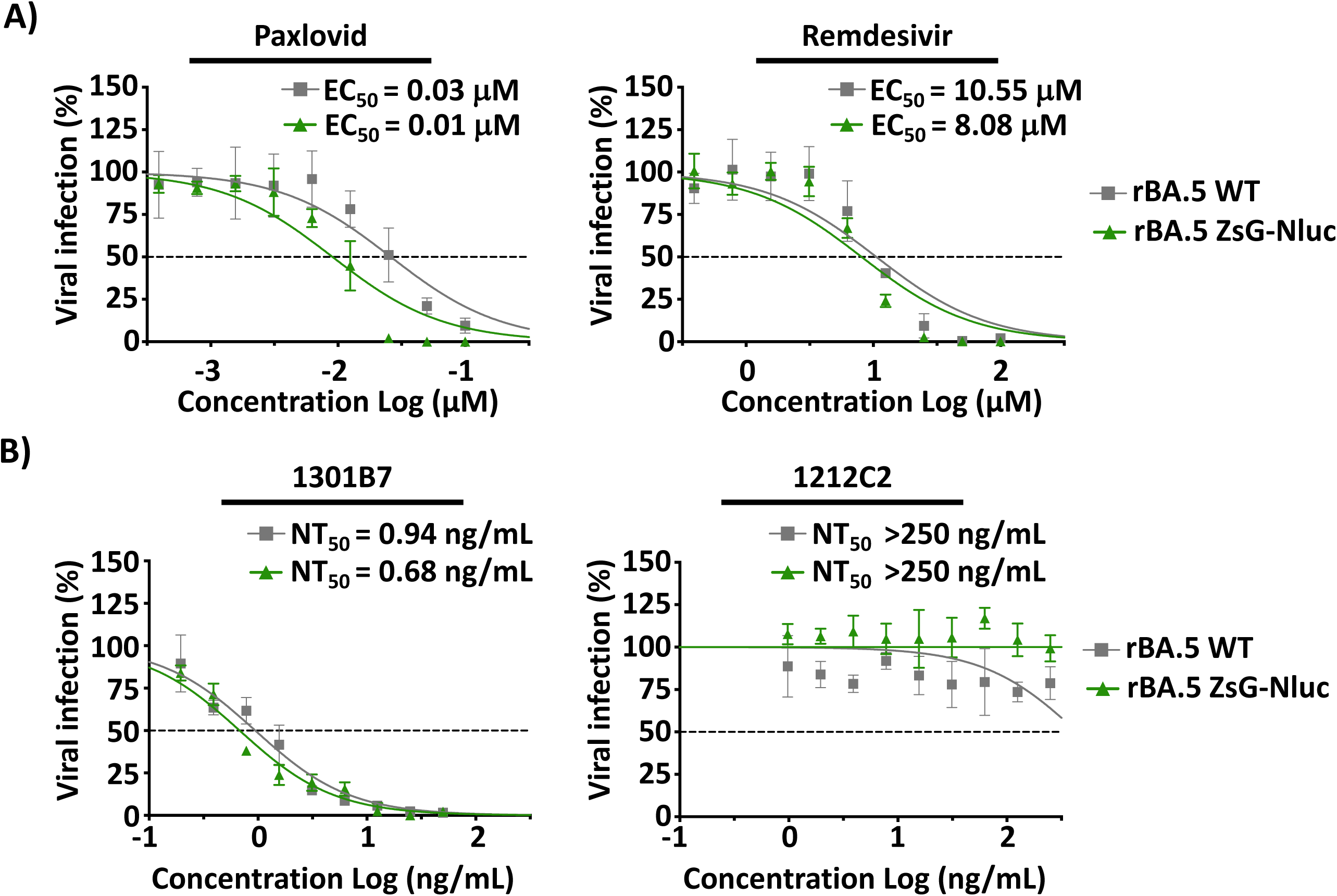
Microneutralization assays to evaluate antivirals and NAbs against rBA.5 ZsG-Nluc. (**A**) Antiviral assay: A549-hACE2 cells were infected (∼100–200 PFU/well) with rBA.5 WT or rBA.5 ZsG-Nluc for 1 h, followed by treatment with 2-fold serial dilutions of Paxlovid (starting concentration of 100 nM) or Remdesivir (starting concentration of 100 µM) in 1% Avicel post-infection media. (**B**) Neutralization assay: A549-hACE2 cells were infected similarly than (**A**) but treated with 2-fold serial dilutions of 1212C2 (starting concentration of 5 µg/mL) or 1301B7 (starting concentration of 1 µg/mL). For rBA.5 ZsG-Nluc (green), viral inhibition was quantified by measuring ZsG expression using a Bioreader 7000 F-z. For rBA.5 WT (gray), cells were stained with the N protein monoclonal antibody 1C7C7 and viral plaques quantified using a Bioreader 7000 F-z.

The majority of NAb therapies have become obsolete in the face of recent SARS-CoV-2 VoC, particularly Omicron, due to the numerous amino acid changes in the receptor binding domain (RBD) of the viral Spike (S) protein (49, 50). However, a reduced set of NAbs isolated from B cells from convalescent individuals targeting a constant region proximal to the hACE2 binding site of the S protein (e.g.1301B7) (34) and those targeting the most conserved S2 domain (e.g. 1249A8) (51) have been shown to effectively neutralize SARS-CoV-2 Omicron strains (34). To explore the feasibility of using rBA.5 ZsG-Nluc to identify NAbs able to neutralize contemporary strains, we performed a neutralization assay with rBA.5 ZsG-Nluc using NAb 1212C2 (42), effective against ancestral WA.1 but not effective against recent Omicron, or using 1301B7 (34), a NAb able to neutralize ancestral and contemporary SARS-CoV-2 strains. Both rBA.5 ZsG-Nluc and rBA.5 WT were efficiently neutralized by 1301B7 with similar NT_50_ (0.68 ng/mL and 0.94 ng/mL, respectively), contrary to 1212C2, which showed no reactivity with Omicron strains (NT_50_ > 250 ng/mL) (**Fig. 3B**). The similarities in susceptibility to antivirals and NAbs with rBA.5 ZsG-Nluc and rBA.5 WT demonstrate the feasibility of using rBA.5 ZsG-Nluc to easily identify and quantify the efficacy of prophylactics and therapeutics *in vitro* by fluorescent or luciferase expression without the need of secondary approaches required to detect the presence of rBA.5 WT in infected wells. More importantly, the use of rBA.5 ZsG-Nluc in these microneutralization assays relies on a more contemporary strain than the original SARS-CoV-2 WA.1.

We next determined the feasibility of using a bireporter fluorescent assay to identify prophylactic or therapeutic candidates against ancestral and Omicron strains, employing our previously described rWA.1 mCherry-Nluc in combination with rBA.5 ZsG-Nluc. As a proof of concept for therapeutic antivirals, we tested Paxlovid and Remdesivir (**Figs. 4A and 4B**). As anticipated, rWA.1 mCherry-Nluc and rBA.5 ZsG-Nluc alone (EC_50_ = 0.018 µM and 0.011 µM, respectively) were inhibited by Paxlovid. Likewise, Remdesivir inhibited both rWA.1 mCherry-Nluc and rBA.5 ZsG-Nluc alone (EC_50_ = 22 µM and 10 µM, respectively) (**Fig. 4A).** In cells infected with both viruses, rWA.1 mCherry-Nluc or rBA.5 ZsG-Nluc were similarly inhibited by Paxlovid (EC_50_ = 0.013 µM and 0.0065 µM, respectively) and Remdesivir (EC_50_ = 15 µM and 5.8 µM, respectively) (**Fig. 4A**). EC_50_ of Paxlovid and Remdesivir against rWA.1 mCherry-Nluc and rBA.5 ZsG-Nluc alone or in combination correlate with levels of mCherry and ZsG expression, respectively, and was observed in images obtained from infected 96-well plates (**Fig. 4B**). Wells treated with the lowest concentration of antiviral (Low) have similar PFU counts to that of untreated infection control images ((-) R_x_), while images from wells treated with EC_50_ values (EC_50_) show half PFU counts. Moreover, images of wells treated with the high concentrations of both antivirals (High) have no PFU counts, comparable to images from mock-infected wells (M).

**Figure 4.**
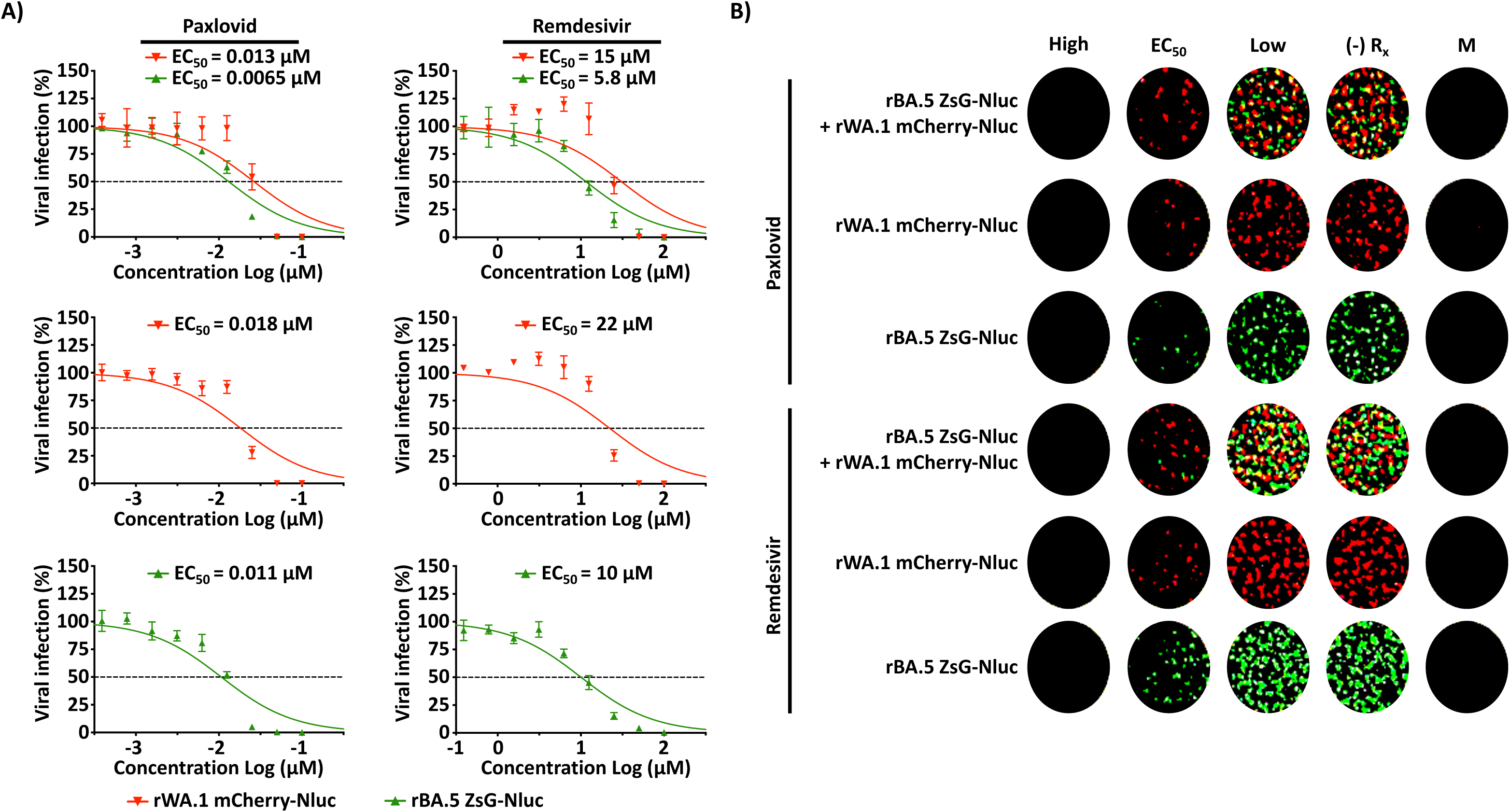
A Bireporter microneutralization assay to evaluate compounds against ancestral and Omicron BA.5. (**A**) A549-hACE2 cells were infected (∼100-200 PFU/well) with rWA.1 mCherry-Nluc (red), rBA.5 ZsG-Nluc (green), or rWA.1 mCherry-Nluc + rBA.5 ZsG-Nluc for 1 h, followed by treatment with 2-fold serial dilutions of Paxlovid (starting concentration of 100 nM) or Remdesivir (starting concentration of 100 µM) in 1% Avicel post-infection media. After fixation, mCherry (red) or ZsG (green) (+) cells were imaged using a Bioreader 7000 F-z. (**B**) Representative images of infections at high, EC_50_, low, no drug (-) Rx, or mock-infected cells (M).

In terms of NAbs, the ancestral rWA.1 mCherry-Nluc was neutralized by 1212C2 (NT_50_ = 1.3 ng/mL), in contrast to rBA.5 ZsG-Nluc that was not neutralized by 1212C2 (NT_50_ > 50 ng/mL) (**Fig. 5A**). Contrarily, the most recent NAb 1301B7 was able to neutralize both rWA.1 mCherry-Nluc and rBA.5 ZsG-Nluc although with different potency (2.4 ng/mL and 0.68 ng/mL, respectively) (**Fig. 5A**). Notably, in A549 cells infected with rWA.1 mCherry-Nluc + rBA.5 ZsG-Nluc, rWA.1 mCherry-Nluc was neutralized by both NAbs 1212C2 and 1301B7 (NT_50_ = 1.6 ng/mL and 3.7 ng/mL, respectively) and rBA.5 ZsG-Nluc was only neutralized by 1301B7 (NT_50_ = 3.1 ng/mL) (**Fig. 5A**). These dose dependent inhibition results with 1212C2 and 1301B7 NAbs correlate with the levels of fluorescent expression seen in the images obtained from a microplate plate reader (**Fig. 5B**). These results demonstrate the feasibility of using ancestral and recent Omicron rSARS-CoV-2 expressing different fluorescent proteins to identify antivirals or NAbs able to inhibit or neutralize, respectively, individually or together original or currently circulating SARS-CoV-2 strains within the same assay.

**Figure 5.**
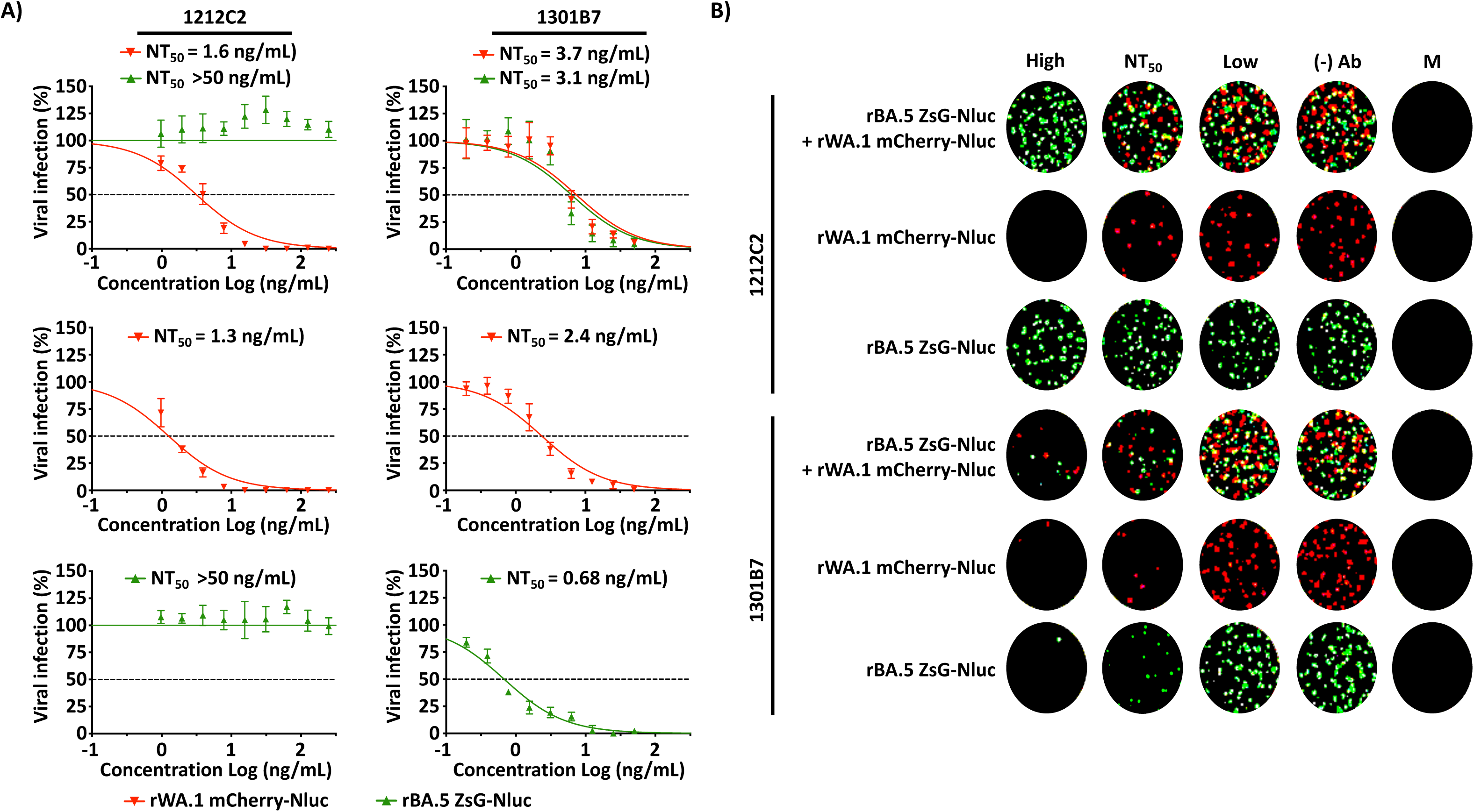
**Bireporter microneutralization assays to evaluate NAbs against ancestral and Omicron BA.5**. (A) rWA.1 mCherry-Nluc (red), rBA.5 ZsG-Nluc (green), or rWA.1 mCherry-Nluc + rBA.5 ZsG-Nluc (∼100-200 PFU/well) were incubated with 2-fold serial dilutions of 1212C2 (starting concentration of 5 µg/mL) or 1301B7 (starting concentration of 1 µg/mL) for 1 h at RT. A549-hACE2 cells were infected with the Ab/virus inoculum for 1 h and overlayed with 1% Avicel post-infection media until fixation and subsequent imaging of mCherry (red) or ZsG (green) (+) cells using a Bioreader 7000 F-z. (B) Representative images of infections at high, NT_50_, low, no Ab (-) Ab, or mock-infected cells (M).

### Genetic stability of rBA.5 ZsG-Nluc

Genetic stability is important for the use of recombinant viruses expressing reporter genes. To assess the genetic stability of rBA.5 ZsG-Nluc, a P2 stock was serially passaged 4 times (P6) in Vero AT cells. The plaque phenotype of P2 and P6 rBA.5 ZsG-Nluc show that all plaques detected by crystal violet staining are also positive for ZsG and Nluc (**Fig. 6A**). Nluc activity in CCSs collected was comparable across all viral passages (P2-P6) (**Fig. 6B**). NGS of rBA.5 ZsG-Nluc P6 identified single nucleotide polymorphisms in >30% of reads at NSP9 (T4158I), NSP13 (M5980I), S (V971G), E (DV14), and N (L13P) and N transcriptional regulatory sequence (TRS) (**Fig. 6C and Supp. Table 1**). However, no mutations were found in the ZsG-Nluc fusion protein, or in the PTV-1 2A sequence (**Fig. 6C and Supp. Table 1**), demonstrating genetic and stability of rBA.5 ZsG-Nluc for at least 6 consecutive passages in Vero AT cells.

**Figure 6.**
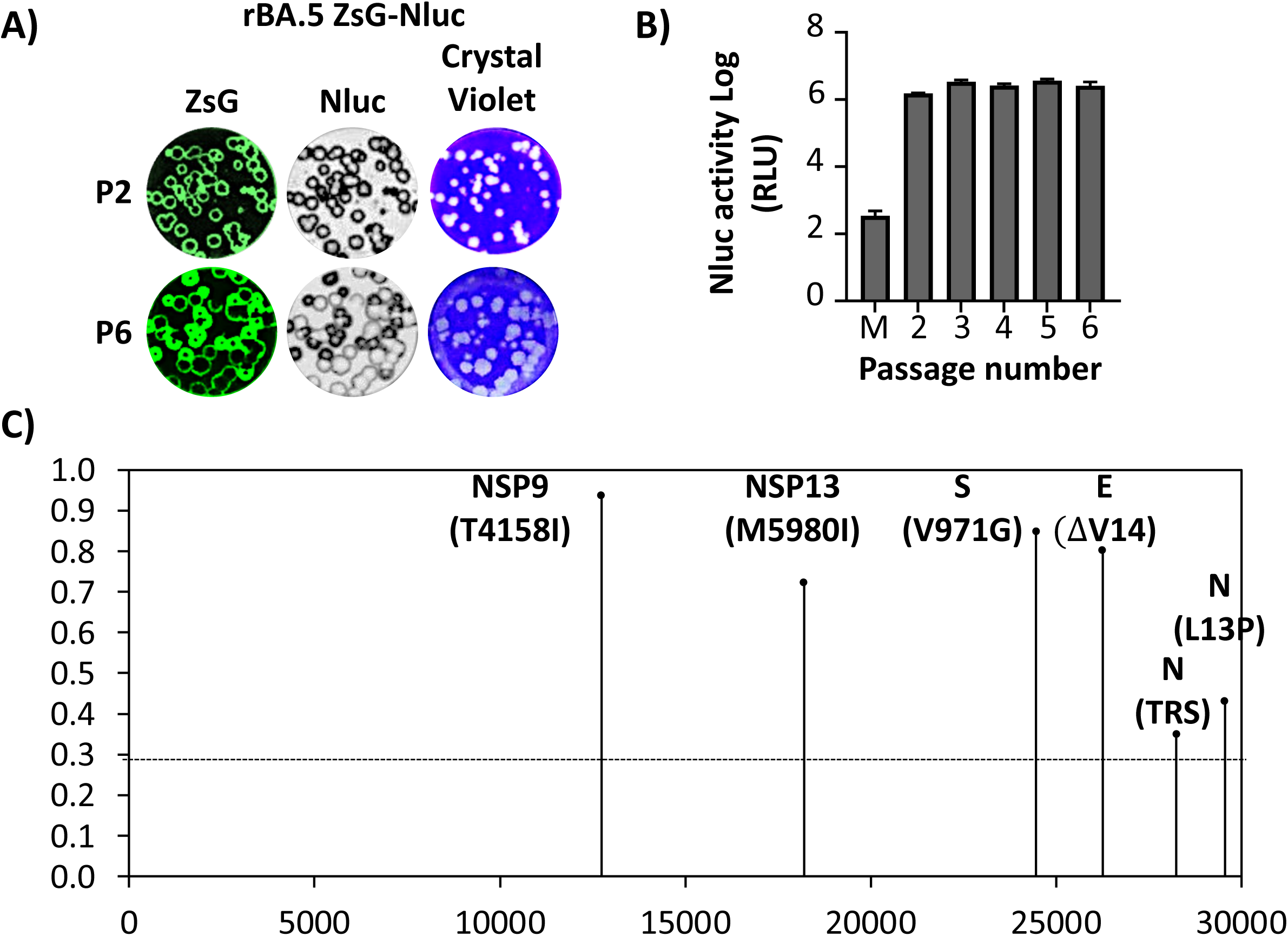
Genetic and phenotypic stability of rBA.5 ZsG-Nluc. Vero AT cells were infected (25-50 PFU/well) with rBA.5 ZsG-Nluc P2 stock and incubated at 37°C for 48 h. At 48 hpi, CCSs were used for quantification of Nluc and subsequent estimation of viral titers for subsequent passages. (**A**) Plaque phenotype of P2 and P6 rBA.5 ZsG-Nluc. (**B**) Nluc activity of different subsequent viral passages. (**C**) RNA from CCS from Vero AT cells infected with rBA.5 ZsG-Nluc P6 was purified by TRIzol and used for NGS genomic analysis.

### Pathogenicity of rBA.5 ZsG-Nluc in K18 hACE2 transgenic mice

A virulent animal model is essential for *in vivo* evaluation of SARS-CoV-2 pathogenicity and assessing the efficacy of candidate vaccines and antivirals (52). Original Omicron strains of SARS-CoV-2 (e.g. BA.1) did not cause morbidity changes in infected K18 hACE2 mice and all animals survived infection with limited virus detected in the lungs (32, 53). We have found that SARS-CoV-2 Omicron BA.5, contrary to its ancestor BA.1, causes high morbidity and mortality, with high levels of viral replication in the lungs of adult (5–8-month-old) male and female K18 hACE2 mice (32). To evaluate if insertion of the ZsG-Nluc in the viral genome or its expression affected pathogenicity or viral replication success, we infected 5–8-month-old male K18 hACE2 mice (n=5) intranasally (i.n.) with 5 × 10^5^ PFU of rBA.5 ZsG-Nluc, a dose we previously showed to result in 100% lethality with rBA.5 WT in similarly aged K18 hACE2 mice. We also included mock-infected mice and mice infected with rBA.5 WT as controls (n=5). Nluc expression in all mice groups were imaged at 1, 2, 4, and 6 dpi using IVIS and followed for 14 days to assess morbidity and mortality (**Fig. 7A**). Over the course of infection until humane endpoints, we were able to detect Nluc expression in mice infected with rBA.5 ZsG-Nluc (**Fig. 7B**). Notably Nluc expression was time dependent, with higher levels of Nluc expression in mice infected with rBA.5 ZsG-Nluc by 6 dpi (**Fig. 7B**). As expected, we could not detect Nluc expression in mice mock-infected or infected with rBA.5 WT (**Fig. 7B**). The bodyweight of rBA.5 ZsG-Nluc- and rBA.5 WT-infected mice steadily decreased over the course of infection (**Fig. 7C**) and all animals succumbed by 6-7 dpi, with no significant differences as determined by Kaplan-Meier curves (**Fig 7D**). We observed moderate body weight loss in mock-infected mice (Mock IVIS). However, this body weight loss was due to anesthesia and retro-orbital injections during the course of the experiment since mock-infected animals that did not undergo IVIS maintained their initial body weight (Mock) (**Fig. 7C**) (54). These results demonstrate the feasibility of using rBA.5 ZsG-Nluc to track systemic viral infections *in vivo*, using IVIS. Moreover, these results suggest that expression of ZsG-Nluc did not significantly impact the pathogenicity of rBA.5 ZsG-Nluc in 5–8-month-old male K18 hACE2 mice.

**Figure 7.**
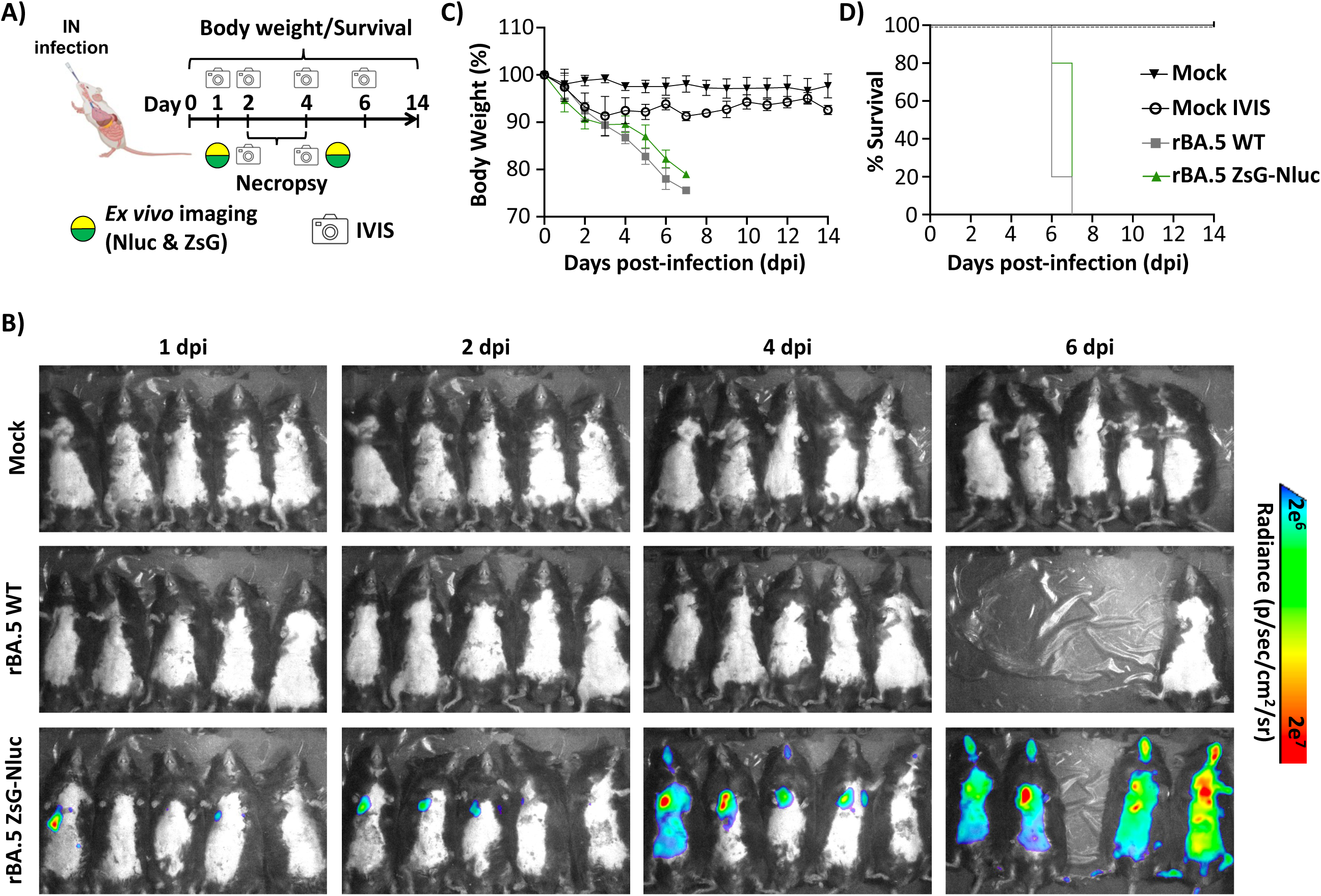
Pathogenicity of rBA.5 ZsG-Nluc in K18 hACE2 transgenic mice. (**A**) Schematic of the *in vivo* experiment. Five- to eight-month-old male K18-hACE2 transgenic mice (n = 5 per group) were i.n. inoculated with 5 × 10⁵ PFU of rBA.5 WT or rBA.5 ZsG-Nluc, or mock-infected. (**B**) At 1, 2, 4, and 6 dpi, mice received retro-orbital injections of Nluc substrate (1:10 in PBS), and whole-body imaging was performed using an IVIS (AMI Spectrum) to visualize Nluc signal. (**C**) Changes in body weight were monitored for 14 dpi. (**D**) Survival was analyzed using Kaplan–Meier curves based on weight loss (%).

### Viral replication of rBA.5 ZsG-Nluc in K18 hACE2 transgenic mice

We next evaluated if rBA.5 ZsG-Nluc was able to replicate to levels comparable to rBA.5 WT, and if Nluc and ZsG expression could be detected *ex vivo* and in the lung tissue from infected mice. To that end, 5–8-month-old male K18 hACE2 mice mock-infected or infected with 5 × 10^5^ PFU of rBA.5 ZsG-Nluc, or rBA.5 WT, were necropsied at 2 (n=4) and 4 (n=4) dpi. Whole body Nluc expression in mock-infected and infected animals was evaluated before euthanasia by IVIS (**Fig. 8A**). As expected, we were able to detect Nluc expression *in vivo* and *ex vivo* in the lungs of mice infected with rBA.5 ZsG-Nluc at 2 and 4 dpi (**Figs. 8A and 8B**). Notably, levels of Nluc expression were higher at 4 than 2 dpi (**Fig. 8B**), results that correspond with the levels of Nluc expression detected in infected animals by IVIS (**Fig. 8A**). Nluc results correlate with ZsG expression levels *ex vivo* in the lungs of mice infected with rBA.5 ZsG-Nluc, with higher levels of ZsG expression at 4 than 2 dpi (**Fig. 8C**). As expected, we could not detect Nluc or ZsG expression in the lungs of mice mock-infected or infected with rBA.5 WT at either 2 or 4 dpi.

**Figure 8.**
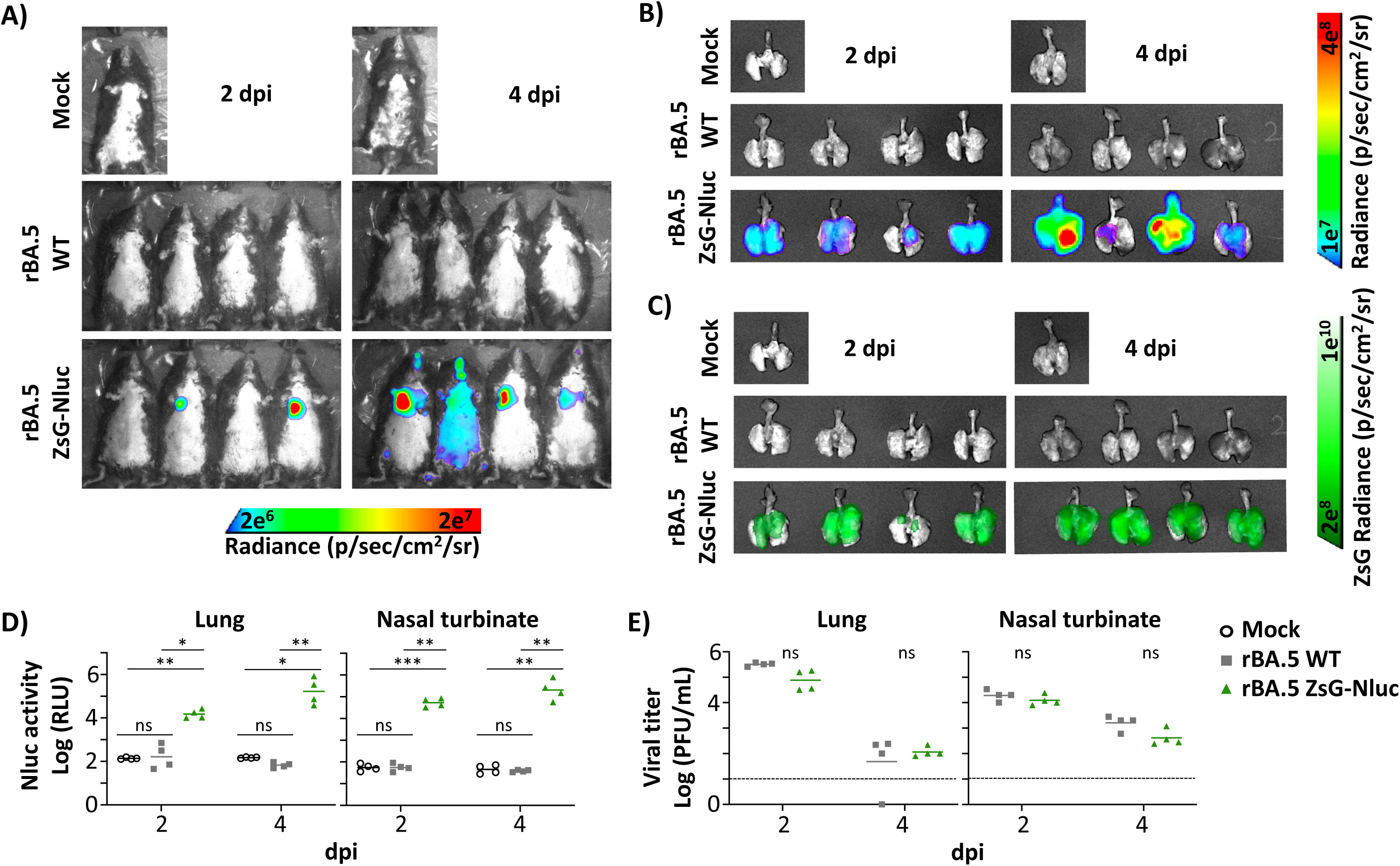
Viral replication of rBA.5 ZsG-Nluc in K18 hACE2 transgenic mice. Five- to eight-month-old male K18-hACE2 transgenic mice (n = 5 per group) were infected i.n. with 5 × 10⁵ PFU of rBA.5 WT or rBA.5 ZsG-Nluc, or mock-infected. At 2 and 4 dpi, mice received retro-orbital injections of Nluc substrate (1:10 in PBS) and subject to whole-body imaging using an IVIS (AMI Spectrum) to visualize Nluc signal (**A**). (**B-C**) *Ex vivo* imaging of lungs from animals in (**A**) were analyzed for Nluc (**B**) and ZsG (**C**) expression using an IVIS. (**D**) Lungs and nasal turbinate tissues were homogenized and analyzed for Nluc signal using a Synergy LX plate reader. (**E**) Viral titers in the same lung and nasal turbinate homogenates were determined by plaque assay. Statistical analysis was performed using one-way ANOVA with Tukey post hoc test. Dotted line indicates the limit of detection (LoD). Data are presented as mean ± SD. ****, P < 0.0001. ***, P < 0.001. ns, non-significant.

After homogenization of lungs and nasal turbinate (NT), homogenates were used to detect Nluc expression (**Fig. 8D**). In both tissue homogenates, Nluc expression was higher at 4 than 2 dpi, correlating with the *in vivo* (**Fig. 8A**) and *ex vivo* (**Fig. 8B**) Nluc and ZsG (**Fig. 8C**) signal intensity. Nluc was undetectable in the lungs or NT of mock-infected or rBA.5 WT-infected mice. Importantly, viral titers in the same lung and NT homogenates showed no statistically significant differences between rBA.5 ZsG-Nluc- and rBA.5 WT-infected mice at 2 or 4 dpi (**Fig. 8E**). These results suggest that rBA.5 ZsG-Nluc has similar pathogenicity and replicates to levels comparable to rBA.5 WT in 5–8-month-old male K18 hACE2. Additionally, we detected systemic presence of rBA.5 ZsG-Nluc in mice *in vivo* based on Nluc activity and *ex vivo* in lung tissues based on ZsG expression. Moreover, we were able to detect Nluc expression in lung and NT tissue homogenates with comparable viral titers for rBA.5 WT and rBA.5 ZsG-Nluc.

## DISCUSSION

Continued circulation of SARS-CoV-2 generates opportunity for the emergence of new VoC, potentially exacerbated by selective immune pressure from previous natural infection or vaccination, or approved antivirals (5, 16, 55). While no mutations conferring drug-resistant to molnupiravir have been reported for SARS-CoV-2 (56) and the prodrug has a high barrier to antiviral resistance (57), molnupiravir remains a 3^rd^ line treatment option for low to moderate COVID-19 with limited efficacy for severe disease in the clinical setting (58, 59). Thus, new prophylactic vaccines and therapeutic antivirals are needed to protect the public from such SARS-CoV-2 emerging VoC. Traditionally, researchers have identified viral infection in cell culture or tissue samples by using antibodies specific to viral proteins, an approach that can be costly and time consuming (60, 61). In addition, natural viral isolates do not allow for tracking viral infections *in vivo*, and multiple cohorts of animals are needed to study viral pathogenesis and replication over the course of an experiment, enhancing the complexity of *in vivo* studies (43, 62–64). The use of recombinant viruses expressing reporter genes has facilitated easy tracking of viral infections *in vitro*, *ex vivo*, and *in vivo* using different fluorescent or luciferase proteins (20, 21, 26, 30, 62, 65). With the advent of new SARS-CoV-2 variants, recovery of recombinant viruses that represent currently circulating strains is required (8, 46, 66). Here, we used a previously described BAC-based reverse genetics system for SARS-CoV-2 BA.5 to generate a bireporter fluorescent (ZsG) and luciferase (Nluc) expressing recombinant virus (rBA.5 ZsG-Nluc) to facilitate *in vitro*, *ex vivo*, and *in vivo* studies with a contemporary viral strain. By inserting the fusion reporter cassette upstream of the gene encoding the viral N protein, which is most abundantly expressed, we detected high levels of reporter expression in cultured cells without affecting stable expression of the bireporter gene up to 6 passages (67–69).

Compared to rBA.5 WT, rBA.5 ZsG-Nluc had similar growth kinetics and plaque phenotype. While plaque size is a factor for determining viral attenuation (70–73), the lack of significant differences in viral replication between rBA.5 ZsG-Nluc and rBA.5 WT suggest a lack of attenuation after insertion of the foreign gene in the viral genome, as previously described with a rSARS-CoV-2 expressing mCherry-Nluc based on the original WA.1 strain (20). Importantly, we demonstrate the feasibility of using rBA.5 ZsG-Nluc for the identification of antivirals and NAbs with EC_50_ and NT_50_, respectively, comparable to those obtained with rBA.5 WT. Moreover, we combined rBA.5 ZsG-Nluc with the previously described rWA.1 mCherry-Nluc to develop a bireporter assay to simultaneously identify antivirals or NAbs able to inhibit or neutralize, respectively, ancestral, Omicron, or both ancestral and Omicron strains simultaneously within the same assay.

*In vivo*, rBA.5 ZsG-Nluc retained the same pathogenicity as rBA.5 WT in aged K18 transgenic hACE2 male mice with no difference in morbidity and mortality, as indicated by Kaplan–Meier curves, or viral titers in lung and NT homogenates. Importantly, live whole body (Nluc) and *ex vivo* (Nluc and ZsG) imaging efficiently detected viral infection in rBA.5 ZsG-Nluc-infected mice. Notably, we also detected Nluc expression in the same tissue homogenates from rBA.5 ZsG-Nluc-infected mice at 2 and 4 dpi, confirming feasibility of using Nluc expression as a valid surrogate readout for viral infection. While our results revealed the efficacy of our bireporter rBA.5 ZsG-Nluc for detection of viral infection in mouse lungs and nasal turbinate, K18-hACE2 mice infected with a natural isolate of Omicron BA.5 have been shown to efficiently replicate in extra-pulmonary organs including the brain, heart, liver, kidney, and colon (74).

However, in our previous studies, and despite causing significant weight loss and mortality, infection with Omicron BA.5, we did not observed any neurological symptoms and we did not detect the presence of the virus, or viral RNA, in the brains of K18-hACE2 mice infected with Omicron BA.5 (32). Future studies using rBA.5 ZsG-Nluc may demonstrate systemic infection in other organs known to be susceptible to Omicron BA.5 infection in K18-hACE2 mice by Nluc (*in vivo* or *ex vivo*) or ZsG (*ex vivo*) expression.

Altogether, we describe the generation and characterization of a bireporter (ZsG-Nluc) rSARS-CoV-2 Omicron BA.5 strain that constitutes a useful tool for *in vitro*, *ex vivo*, and *in vivo* studies, including the identification and characterization of therapeutics against a recent SARS-CoV-2 variant. This dual-reporter strain can be used alone, or in combination with the ancestral WA.1 strain in a bireporter assay that allows the simultaneous identification of therapeutics against both, original and more recently emerged SARS-CoV-2 strains within a single assay.

## MATERIALS AND METHODS

### Biosafety

All *in vitro*, *ex vivo*, and *in vivo* studies involving SARS-CoV-2 were carried out at biosafety level 3 (BSL-3) and animal BSL-3 (ABSL-3) facilities at Texas Biomedical Research Institute. Studies were approved by the Institutional Biosafety Committee (IBC 20-004) and the Institutional Animal Care and Use Committee (IACUC 1718) at Texas Biomedical Research Institute.

### Cells

Vero cells expressing human angiotensin converting enzyme 2 (hACE2) and transmembrane serine protease 2 (TMPRSS2) (Vero AT) and A549 human epithelial lung cells expressing hACE2 (A549 hACE2) were obtained from BEI Resources (NR-54970 and NR-53821, respectively), and grown in Dulbecco’s modified Eagle’s medium (DMEM) supplemented with 10% fetal bovine serum (FBS) and 1% PSG (penicillin,100 units/ml; streptomycin 100 μg/ml; L-Glutamine, 2 mM) at 37°C in a 5% CO_2_ cell culture incubator.

### Plasmids and viruses

Recombinant (r)SARS-CoV-2 BA.5 wild-type (rBA.5 WT) and rSARS-CoV-2 BA.5 expressing ZsGreen (ZsG) and NanoLuc (Nluc) (rBA.5 ZsG-Nluc) were constructed and rescued as previously described (31). Briefly, rBA.5 ZsG-Nluc contains a ZsG-Nluc fusion protein and a downstream porcine teschovirus-1 (PTV-1) 2A autoproteolytic cleavage site (ATNFSLLKQAGDVEENPGP) inserted upstream of the nucleocapsid (N) open reading frame (ORF) in the SARS-CoV-2 BA.5 viral genome (20). Plasmid construct was confirmed by DNA next generation sequencing (Plasmidsaurus). The rBA.5 WT and rBA.5 ZsG-Nluc viruses were rescued as previously described by transfecting Vero AT cells with 2 µg of the corresponding bacterial artificial chromosomes (BACs) using lipofectamine 2000 (LPF2000) (31). Rescued viruses were plaque purified and amplified in Vero AT cells to generate viral stocks as previously described (32). Viral stocks were aliquoted and stored at −80°C until further use. Viral titers were determined by standard plaque assay in Vero AT cells. The rSARS-CoV-2 Washington 1 (WA.1) strain expressing a fusion of mCherry-Nluc using the same PTV-1 2A approach from the locus of the N viral protein (rWA.1 mCherry-Nluc) was previously described (20).

### Fluorescent microscopy

Fluorescent expression in Vero AT cells infected with rBA.5 WT or rBA.5 ZsG-Nluc was visualized as previously described (20). Briefly, Vero AT cells (12-well plate format, 2.5 x10^5^ cells/well, triplicates) were mock (PBS)-infected or infected with rBA.5 WT or rBA.5 ZsG-Nluc at a multiplicity of infection (MOI) of 1 in infection media (2% FBS 1% PSG DMEM) for 1 h at room temperature (RT). After viral adsorption, cells were washed 3X with Phosphate Buffered Saline (PBS), overlayed with infection media and incubated at 37°C in a 5% CO_2_ cell culture incubator. At 12 hours post-infection (hpi), cells were fixed in 10% neutral buffered formalin for 24 h (33). Cells were then permeabilized with 0.5% Triton X-100 in PBS for 10 minutes at RT. Cells were blocked with 2.5% bovine serum albumin (BSA) in PBS at 37°C for 1 h and incubated with a mouse monoclonal antibody (MAb) against the viral N protein (1C7C7) for 1 h at 37°C (34, 35). Cells were then washed 3X with PBS and incubated with an anti-mouse TxRed conjugated polyclonal antibody for 1 h at 37°C. After 3X washes with PBS, cells were stained with 4′,6-diamidino-2-phenylindole (DAPI) for 30 minutes at RT. Cells were next visualized under an EVOS M5000 fluorescent microscope using GFP, TxRed, and DAPI filters.

### Viral growth kinetics

Viral growth kinetics were conducted in Vero AT cells (6-well plate format, 10^6^ cells/well, triplicates). Briefly, cell monolayers were infected at MOI of 0.001. After 1h of viral adsorption at RT, cells were washed 3X with PBS, overlayed with infectious media and incubated at 37°C in a 5% CO_2_ cell culture incubator. At 12, 24, 48, and 72 hpi, cells were imaged using an EVOS M5000 fluorescent microscope and viral titers in the cell culture supernatants (CCSs) were determined by plaque assay as previously described (20). Nluc activity in the CCSs was quantified using the Promega Nluc assay kit and a PheraStar microplate reader. Mean value and standard deviation (SD) were calculated using Microsoft Excel Software.

### Plaque assays

Confluent monolayers of Vero AT cells (6-well plate format, 10^6^ cells/well) were infected with 10-fold serial dilutions of rBA.5 WT or rBA.5 ZsG-Nluc. After 1 h viral adsorption, infectious virus was replaced by semi solid DMEM/Agar infection media, and the plates were incubated at 37°C in a 5% CO_2_ cell culture incubator. At 72 hpi, plates were fixed with 10% neutral buffered formalin for 24 h. For visualization of ZsG expression, the agar overlay was removed, and cells were washed 3X with PBS, incubated in 1 mL of PBS and imaged using an Alexa 488 filter on a BioRad ChemiDoc MP imaging system (Chemidoc). Subsequently, for Nluc visualization, cell monolayers were incubated with Nluc substrate (1:200 dilution in PBS) and imaged using a Chemidoc. Following Nluc visualization, cell monolayers were incubated with 1 mL of 0.1% crystal violet solution at RT for 15 minutes (36). Cells were washed 3X with PBS and dried to be imaged using an Epson Perfection V600 Photo Scanner.

### Genetic stability of rBA.5 ZsG-Nluc *in vitro*

Vero AT cells (6-well plate format, 10^6^ cells/well, triplicates) were infected at MOI of 0.001 with rBA.5 ZsG-Nluc passage 2 (P2) stock and incubated at 37°C in a 5% CO_2_ cell culture incubator until ∼90% cytopathic effect (CPE) was observed (∼48 hpi). To normalize viral titer between passages, original P2 viral stock or infected CCSs were incubated with 1:1 volume of Nluc substrate diluted in PBS (1:100). Nluc activity was measured using a PheraStar instrument. Nluc values in each viral passage were normalized to Nluc values in the P2 viral stock. After 4 consecutive passages in Vero AT cells, viral RNA from P2 and P6 CCSs were extracted using TRIzol LS reagent (Invitrogen) using manufacturer’s instructions and analyzed via whole genome sequencing using next-generation sequencing (NGS) MinION (Oxford Nanopore Technologies, or ONT). Sample libraries were assayed with the Native Barcoding Kit 24 V14 (SQK-NBD114.24, ONT) and ran on R10.4.1. Flow Cells (FLO-MIN114, ONT) per manufacturer’s instructions. The raw read length stats were created with n50 v1.7.0 and trimmed using nanoq v0.10.0 (37). Bases removed from downstream analyses included the first and last 25 bp of each read, those with PHRED quality scores < 7, and reads < 500 bp after trimming. Filtered reads were mapped to the rBA.5 ZsG-Nluc reference sequence using minimap2 v2.28-r1209 (38) and the ‘-x map-ont’ option for mapping long error prone nanopore reads. Mapping rates were calculated with SAMtools flagstat v1.21 (39). Coverage across each genome was estimated with MosDepth v0.3.10 (40). Indel quality scores were re-assessed, and variants were called using LoFreq v2.1.5 (41). Depth of coverage (--max-depth) for variant calling was limited at 10,000X and default filters applied by LoFreq were removed. Rather, variants in < 30% of reads or regions of the genome that contained less than 100x read depth were removed.

### Microneutralization assays

Antiviral microneutralization assays were conducted as previously described (20). Briefly, confluent monolayers of A549 hACE2 cells (96-well plate format, 5×10^4^ cells/well, quadruplicates) were infected with 100-200 plaque-forming units (PFU) of rBA.5 WT or rBA.5 ZsG-Nluc and incubated for 1 h in a 5% CO_2_ cell culture incubator. After viral adsorption, cells were then overlayed with 1% Avicel in infection media containing 2-fold serial dilutions of Paxlovid or Remdesivir (starting drug concentration of 100 nM and 100 µM, respectively) and incubated at 37°C in a 5% CO_2_ cell culture incubator. At 24 hpi, plates were fixed in 10% neutral buffered formalin for 24 h. Cells were then washed 3X with PBS and underwent immunostaining, or overlayed with 300 µL of PBS if used for fluorescent imaging. For immunostaining, cells were permeabilized with 0.5% triton X-100 in PBS for 15 minutes at RT, washed 3X with PBS, blocked with 2.5% BSA in PBS for 1 h at 37°C, and incubated with the mouse anti-N 1C7C7 MAb in 1% BSA in PBS for 1 h at 37°C. After incubation with the primary antibody, cells were washed 3X with PBS and incubated with an anti-mouse secondary VectaStain Ab for 30 minutes at 37°C. After incubation with the secondary antibody, cells were washed 3X with PBS and incubated with ABC reagent for 30 minutes at 37°C. After, cells were washed 3X with PBS and incubated with DAB reagent for 5 minutes at RT. Immunostained plaques or fluorescent plaques were then imaged in a Bioreader 7000 F-z instrument. Microneutralization curves were created using Microsoft Excel and GraphPad Prism software. The average of the quadruplicate wells was used to calculate the SD of antiviral activity using Microsoft Excel Software. The half-maximal effective concentration (EC_50_) was calculated using a sigmoidal dose-response curve (GraphPad Prism software).

NAb assays were conducted as previously described (21). Briefly, 100-200 PFU of rBA.5 WT or rBA.5 ZsG-Nluc were incubated with infection media containing 2-fold serial dilutions of the previously described 1212C2 or 1301B7 human MAbs (34, 42) (starting concentrations of 5 µg/mL or 1 µg/mL, respectively) for 1 h at 37°C in a 5% CO_2_ cell culture incubator. Next, 96-well plates of A549 hACE2 cells (4 x 10^4^ cells/well, quadruplicates) were (+/-) infected with the viral/antibody mixture for 1 h at RT. After 1h viral absorption, infectious media was removed and cells were incubated in infection media for 24 h at 37°C in a 5% CO_2_ cell culture incubator. At 24 hpi, plates were fixed in 10% neutral buffered formalin for 24 h. Cells were then washed 3X with PBS and immunostained or overlayed with 300 µL of PBS for fluorescent imaging as described above. Fluorescent plaques were then imaged in a Bioreader 7000 F-z instrument and underwent analysis as described above to determine the 50% neutralizing titer (NT_50_) of the antibodies. NT_50_ were calculated using a sigmoidal dose-response curve (GraphPad Prism software).

### Bireporter microneutralization assays

Bireporter antiviral and NAb microneutralization assays were conducted as previously described (20, 21). Viral stocks were titrated in A549 hACE2 cells using 96-well plates (4 x 10^4^ cells/well, quadruplicates) using 2-fold serial dilutions of rWA.1 mCherry-Nluc, rBA.5 ZsG-Nluc, or rWA.1 mCherry-Nluc + rBA.5 ZsG-Nluc. After 1h viral absorption at RT, infectious media was removed and cells were incubated in infection media for 24 h at 37°C in a 5% CO_2_ cell culture incubator. At 24 hpi, plates were fixed in 10% neutral buffered formalin for 24 h. Cells were then washed 3X with PBS and overlayed with 300 µL of PBS for fluorescent imaging as described above using a Bioreader 7000 F-z instrument. Viral inoculum that resulted in 100-200 fluorescent plaques of rWA.1 mCherry-Nluc, rBA.5 ZsG-Nluc, or rWA.1 mCherry-Nluc + rBA.5 ZsG-Nluc became standard volumes for use in a single well of infection in the bireporter assays using antivirals and NAbs. For antiviral and NAb assays, drug and NAb concentrations followed the same starting concentrations as described above. The EC_50_ and NT_50_ of antivirals and NAbs, respectively, in cells co-infected with rWA.1 mCherry-Nluc + rBA.5 ZsG-Nluc were normalized to those using single viral infections and calculated using a sigmoidal dose-response curve (GraphPad Prism software).

### Mouse experiments

For whole body *in vivo* imaging, body weight changes, and survival, 5–8-month-old male Keratin-18 human angiotensin converting enzyme-2 (K18 hACE2) mice (43) (n=5) were purchased from The Jackson Laboratory and housed under specific pathogen-free conditions at Texas Biomedical Research Institute. For viral infection, mice were anesthetized intraperitoneally (i.p.) with a mixture of Ketamine (100 mg/mL) and Xylazine (20 mg/mL) (44, 45). Mice were intranasally (i.n.) inoculated with 50 µL of either rBA.5 WT or rBA.5 ZsG-Nluc (5 X 10^5^ PFU/mouse) (32). *In vivo* Nluc imaging was performed using the AMI IVIS Spectrum multispectral imaging system. At 1-, 2-, 4-, and 6-days post infection (dpi), mice were anesthetized and injected retro-orbitally with 100 µL of Nano-Glo luciferase substrate (Promega)f in PBS (1:10 dilution). Immediately following injection, bioluminescence was measured using an AMI Spectrum IVIS. Bioluminescence data was acquired and analyzed using the Aura program (AMI Spectrum). Morbidity, as indicated by changes in body weight, and mortality, as assessed by survival rate, were monitored over 14 days. Mice that lost 20% or more of their initial body weight were considered to have reached the experimental endpoint and were humanely euthanized. Survival data was analyzed using Kaplan-Meier curves (20, 21).

For viral titrations, 5–8-month-old male K18 hACE2 mice (n=8) were similarly infected and underwent whole body IVIS imaging at 2 and 4 dpi as described above. For *ex vivo* imaging, lungs were collected following euthanasia with i.p. injections of 100 mL of Fatal Plus solution. After imaging, lungs and nasal turbinate (NT) were homogenized in 1 mL of PBS using a Precellys tissue homogenizer (Bertin Instruments) for 20 seconds at 7,000 rpm. Tissue homogenates were then clarified via centrifugation at 12,000 x g at 4°C for 5 minutes, and the tissue supernatants were collected for titration by plaque assay as described above. Tissue homogenates were also used to assess levels of Nluc using the Nano-Glo luciferase substrate (1:100 in PBS) and a BioTek Synergy LX microplate reader.

### Statistical analyses

All graphs, calculations, and statistical analyses were performed using GraphPad Prism software version 9.5.1 (GraphPad Software, LLC, USA) and Microsoft Excel version 16.95.4.

## Acknowledgments

We thank BEI resources for providing the Vero AT and A549 hACE2 cells.

## Authors Contributions

**E.M.C.:** Conceptualization, Methodology, Investigation, Data curation, Formal analysis, Visualization, Writing – original draft;

**R.S.B.:** Investigation, Methodology, Writing – review & editing;

**C.Y.:** Resources, Investigation, Methodology, Validation, Writing – review & editing; **B.I.:** Investigation, Methodology – aged mouse model, Writing – review & editing; **S.E.:** Investigation, Methodology – aged mouse model, Writing – review & editing;

**A.C.:** Formal analysis, Data curation, Validation, Writing – review & editing; **M.R.W.:** Investigation, Methodology – NAb discovery, Funding acquisition, Writing – review & editing;

**J.J.K.:** Investigation, Methodology– NAb discovery, Funding acquisition, Writing – review & editing;

**R.K.P.:** Investigation, Methodology, Funding acquisition, Writing – review & editing;

**A.G.-S.:** Supervision, Funding acquisition, Writing – review & editing;

**A.M.:** Investigation, Methodology, Writing – review & editing;

**H.C.A.:** Investigation, Methodology – aged mouse model, Writing – review & editing; **L.M.-S.:** Conceptualization, Supervision, Funding acquisition, Project administration, Writing – review & editing.

All authors reviewed and approved the final manuscript.

---

**Supplementary table 1.** Amino acid changes in rBA.5 ZsG-Nluc P6.

| **Change** | **Coverage** | **Polymorphism Type** | **Allele Frequency** | **Variant P-Value (approximate)** | **Amino Acid Change** |
| --- | --- | --- | --- | --- | --- |
| C -> T | 566 | Single nucleotide (transition) | 0.94 | 0 | NSP9 (T4158I) |
| G -> A | 1,317 | Single nucleotide (transition) | 0.72 | 0 | NSP13 (M5980I) |
| T -> G | 1,341 | Single nucleotide (transversion) | 0.85 | 0 | S (V971G) |
| -GTT | 227 | Deletion | 0.80 | 9.2E-233 | E (ΔV14) |
| T -> A | 57 | Single nucleotide (transversion) | 0.35 | 6E-20 | N (TRS) |
| T -> C | 37 | Single nucleotide (transition) | 0.43 | 1.8E-27 | N (L13P) |

